# Netrin-1 regulates colorectal cancer stem cell self-renewal via a TFF3 dependent paracrine survival mechanism

**DOI:** 10.1101/2025.04.28.650943

**Authors:** Morgan Brisset, Kristina Radkova, Andrea Paradisi, Mélodie Grandin, Lea Stephan, Robin Wagner, Cyril Degletagne, Fabien Luiggi, Lisa Frydman, Alexander G. Heriot, Corina Behrenbruch, Tamara Vu, Patrick Mehlen, Frédéric Hollande

## Abstract

**Background:** Metastatic colorectal cancer (mCRC) is associated with high recurrence rates and resistance to conventional treatments, largely driven by cancer stem cells (CSCs) that contribute to tumor progression and therapeutic evasion. This study aims to investigate the role of netrin-1 and its dependence receptor UNC5B in regulating CSC self-renewal in mCRC and explore their potential as therapeutic targets.

**Methods:** We used patient-derived liver metastasis organoids (PDOs) to examine the effects of netrin-1 on CSC self-renewal. The role of UNC5B was evaluated by silencing its expression using CRISPR and assessing the impact on CSC apoptosis in response to an anti-netrin-1 blocking antibody (NP137) using extreme limiting dilution assays (ELDAs). Single-cell RNA sequencing was employed to explore the molecular mechanisms behind netrin-1/UNC5B regulation of CSC fate. Clinical data from a patient with mCRC were used to validate the findings.

**Results:** Netrin-1 promoted CSC self-renewal by inhibiting apoptosis, a process reversed by NP137. UNC5B was identified as the primary receptor mediating this effect, as its silencing eliminated Netrin-1-induced self-renewal. Trefoil Factor 3 (TFF3), secreted by UNC5B-expressing cells, plays a key role in netrin-1-induced CSC self-renewal. Clinical trial data from a patient with mCRC showed a reduction in TFF3 and stemness genes expression after treatment with NP137. Furthermore, combining NP137 with FOLFOX chemotherapy enhanced cell death and inhibited tumor growth in PDO xenograft models.

**Conclusion:** This study identifies the netrin-1/UNC5B/TFF3 axis as a critical regulator of CSC self-renewal in mCRC and suggests that targeting this pathway with NP137, in combination with chemotherapy, could provide a promising therapeutic approach for mCRC patients.

## Introduction

Colorectal cancer (CRC) remains a leading cause of cancer-related deaths worldwide, with particularly poor prognosis once metastasized (mCRC), where five-year survival rates drop below 13%^1–3^. One of the primary challenges in treating mCRC is the resistance to chemo- and targeted therapies, a resistance driven in part by cancer stem cells (CSCs)^4–6^. These CSCs possess self-renewal and chemoresistant capabilities, making them a key therapeutic target in the search for more effective and longer-lasting treatments^7,8^.

Netrin-1, a protein originally identified as a guidance cue in neural development, has emerged as a key regulator in cancer biology through its interaction with dependence receptors, such as UNC5B^9–12^. Dependence receptors induce apoptosis in the absence of their ligand, such as netrin-1, making them critical players in the balance between cell survival and death^13^. In several solid tumors, netrin-1 was recently shown to promote epithelial-mesenchymal transition (EMT), a process that enhances cancer cell migration, invasion, and metastasis. Inhibition of netrin-1 has been shown to induce cell death, block EMT and tumor progression in clinical and preclinical settings^14,15^.

In this study, we leverage patient-derived metastatic organoid models to demonstrate that netrin-1 promotes the self-renewal capabilities of mCRC CSCs. We further identify that this effect is mediated by UNC5B expressing cells, which respond to netrin-1 binding by secreting Trefoil Factor 3 (TFF3), a stemness-enhancing factor^16,17^. Our findings highlight the potential of targeting the Netrin-1/UNC5B/TFF3 axis to disrupt CSC-driven resistance in mCRC, opening new avenues for therapeutic intervention.

## Material and Methods

### Patient-derived organoid (PDOs)

Patients with CRC were identified prospectively from January 2016 to June 2016. Colorectal liver metastases and, when possible, matched healthy hepatic tissue and primary tumor were collected from patients undergoing treatment at Peter MacCallum Cancer Centre, The Royal Melbourne Hospital, St Vincent’s Hospital, St Vincent’s Private Hospital or Morwell Hospital by Dr Corina Behrenbruch. Tumor samples of approximately 0.25-1cm^3^ were collected and used for the establishment of patient-derived organoids and FFPE blocks as well as RNA sequencing studies. The study leading to the generation of these organoid cell lines was approved by the Peter MacCallum Cancer Centre Human Research Ethics Committee (HREC/15/PMCC/112, project #15/169) and written consent was obtained from each patient according to National Health and Medical Research Council guidelines.

Organoids were grown inside a Growth-factor reduced Matrigel matrix (Corning, Cat #356231) forming a dome at the bottom of the well. After completion of Matrigel polymerization, complete organoid growth medium composed of advanced DMEM/F12 (#2634010, Thermo Fisher Scientific) supplemented with Glutamax (2 mM; Life Technologies, #35050-061), penicillin-streptomycin (10.00 U/mL; Life Technologies, #15140-122), recombinant human EGF (50 ng/mL, Miltenti Biotec, #130-097-749), Leu [15] human Gastrin I (1 μg/mL, Merck, # G9145), N-acetyl cysteine (1 mM, Merck, #A9165), B27 (2X; Life Technologies, #17504044), A83-01 (ALK inhibitor; 500 nM, Merck, #SML0788), SB202190 (p38/MAPK inhibitor; 10 nM, Miltenyi Biotec, #130-106-275) and YP-27632 (ROCK inhibitor; 10 μM, Abcam, #120129) is added. To correct for variation in protein concentration between Matrigel batches, all Matrigel was prediluted to a concentration of 4.35mg/ml with DMEM/-F12.

### CRISPR PDOS

UNC5B sgRNA sequences were derived from the Brunello human CRISPR knockout pooled library^18^. Linearisation of the human FU-Cas9Cherry expression plasmid (Addgene, Ref. #70182) was performed using the Esp□I (BsmBI) restriction enzyme (Thermo Fisher Scientific, Ref. #ER0451). Transfection was made using HEK293T cells and a mix including the packaging vectors (pMDLg/pRRE - Addgene #12251, pRSV-Rev - Addgene #12253 and pMD2.G/VSVg - Addgene #12259) and sgRNA-expressing DNA (1.4µg of the corresponding target-specific sgRNA or 1.4ug of sgScramble used as control)

To edit our organoid cell lines using CRISPR, cells were first transduced with Cas9 and with sgRNA viruses. Organoids were then amplified and sorted using flow cytometry using mCherry as a marker of transduction of the Cas9-plasmid. For sgRNA transductions, an identical protocol was performed on the Cas9-mCh+ cells. After transduction, the Cas9-mCh+/sgRNA-GFP+ double-positive population was sorted by FACS and amplified. CRISPR editing in each monoclonal population was validated by TIDE analysis and western blot.

### Western blots

Whole-cell protein lysate was obtained by suspending cell pellets in Laemli lysis buffer (60 mM Tris-HCl, 10% v/v glycerol, 2% w/v SDS, 2% v/v β-mercaptoethanol) at room temperature for 10 minutes, followed by boiling the samples at 90°C for 10 minutes. Protein concentrations were determined using the Pierce BCA protein assay kit (Thermo Fisher Scientific; 23227). Anti-UNC5B (Cell Signalling; 13851) and anti Netrin-1 (Abcam; ab126729) were used to detect respectively the UNC5B receptor and Netrin-1.

### Quantitative Polymerase Chain Reaction

Total RNAs were extracted utilizing the NucleoSpin® RNA Plus Kit (Macherey Nagel, Düren, Germany) by the manufacturer’s protocol. RT-PCR reactions were conducted using the PrimeScript RT Reagent Kit (Takara Bio Europe, Saint-Germain-en-Laye, France). A total of 500 ng of total RNA underwent reverse transcription with the following program: 37°C for 15 minutes and 85°C for 5 seconds.

For expression studies, the target transcripts were amplified in a LightCycler® 2.0 apparatus (Roche Applied Science, Indianapolis, IN, USA) using the Premix Ex Taq probe qPCR Kit (Takara Bio Europe) following the manufacturer’s instructions. The expression of target genes was normalized to TATA-binding protein (TBP) which served as the housekeeping gene The quantity of target transcripts, normalized to the housekeeping gene, was determined using the comparative CT method: CT = CT(a target gene)−CT(a reference gene). ΔΔCT = ΔCT(a target sample)−ΔCT(a reference sample) = (CTD − CTB)−(CTC − CTA). Primer sequences: NTN1 : Rev : 5’-GGC ATG CAG GTT GCA; For: 5’ GCA AGC CCT TCC ACT; TBP : Rev: 5’-CAC ACG CCA AGA AAC; For: 5’-CGG CTG TTT AAC TTC; UN5B : Rev: 5’-CCA AGT ATC CGC CCA; For: 5’-GAC ACG CCT GTA GCA; TFF3 : For : 5’-TGG GCC TGT CTG CAA ACCA; Rev : GCT TGA AAC ACC AAG GCA CTCC. TFF3 : For : 5’-TGG GCC TGT CTG CAA ACCA; Rev : GCT TGA AAC ACC AAG GCA CTCC.

### Flow cytometry

For all flow cytometry analysis and cell sorting, single-cell suspensions were resuspended in FACS Buffer (2% FBS, 0.2% EDTA in PBS). Raw cytometry data was analysed using FlowJo software (Tree Star Incorporated, v 10.8.1). Cas9-mCh+/sgRNA-GFP+ double-positive population was analysed by flow cytometry. The sorting was done by the Flow Cytometry facility of the Peter MacCallum Cancer Center and was performed on a FACSAriaTM Fusion cytometer.

For AnnexinV and propidium iodide (PI) double staining, cells were then harvested and dissociated into single cells as previously described, washed with PBS and incubated for 10 minutes in Anti-CD133 (Prominin-1) Antibody (MAB4399-I, MilliporeSigma). Cells were then washed and resuspended in Annexin V buffer containing allophycocyanin (APC) conjugated to Annexin V and PI (BioLegend, San Diego, CA, USA). Annexin V-positive cells were then evaluated by fluorescence-activated cell sorting on FACSCanto II (BD Biosciences, San Jose, CA, USA).

### Extreme limiting dilution Assays (ELDAs)

The Extreme Limiting Dilution Assay is a statistical method commonly used in stem cell biology and cancer research. This assay helps estimate the frequency of functional cells with a particular activity within a heterogeneous cell population. It involves diluting the cells to such an extent that each well, on average, contains either one functional cell or none. By observing the proportion of wells exhibiting the desired activity, we can calculate the frequency of the functional cells in the original population. This assay is particularly useful for studying rare cell populations, such as cancer stem cells, where traditional methods might be less sensitive. ELDA was employed to assess the self-renewal capacity of PDO cell lines. The protocol to perform an ELDA on our PDO cell lines was adapted from Hu, Y. et al article: “ELDA: Extreme limiting dilution analysis for comparing depleted and enriched populations in stem cell and other assays ^19^*”.* Cells were then incubated in media containing either recombinant Netrin-1 (100 ng/mL), NP001 isotypic control (10 ug/mL) or NP137 neutralizing antibody (10 ug/mL) for 72 h. For ELDAs evaluating the effect of the caspase inhibitor Z-VAD(OH)-FMK on NP137 effect, Z-VAD(OH)-FMK (Ref: 161401-82-7, Selleck Chemicals) was supplemented to the organoid medium at 50ug/mL. For ELDAs evaluating the effect of NP137 on post-FOLFOX treatment self-renewal, FOLFOX was added to a final concentration of 1 μM oxaliplatin + 1 μM 5-Fluorouracil. Cells were seeded at the following densities: 250, 125, 25, 5 and 1 cell(s) per well, covering the dynamic range of the sphere-forming frequency of our organoid cell lines. Each of the four treatment groups had 10 to 16 replicates for every dilution, depending on the experiment. Experiments assessing the impact of FOLFOX, FOLFOX treatment was performed before seeding the ELDA but not maintained after single-cell seeding. After 12 days in culture, the assessment of organoid formation scoring in each well is based on the binary presence or absence of organoid(s). The fraction of responsive cells at each density was then subjected to analysis using the ELDA tool provided by the Bioinformatics Division of the Walter and Eliza Hall Institute of Medical Research (http://bioinf.wehi.edu.au/software/elda/).

In individual experiments, pairwise differences in stem cell frequency among treatment groups were evaluated using a chi-square test. For aggregated data, statistical analysis was conducted using one-way, unpaired ANOVA, followed by the Bonferroni correction for multiple comparisons.

### Single-cell RNA-seq

Organoid cell lines were seeded onto 48-well plates at a density of 50% confluence. Cells were then incubated in media containing either NP001 isotypic control (10 ug/mL) or NP137 neutralising antibody (10 ug/mL) for 72 h. Organoids were dissociated using TrypLe for 15 minutes at 37°C. Then, organoids were mechanically pipetted until a single-cell suspension was achieved) and passed through a 40 μm nylon mesh cell strainer to exclude large aggregates. The enumeration of viable cells was conducted using a Luna-FL Dual fluorescence cell counter (Logos Biosystems). Cells were loaded on Chip G and run on chromium iX to reach 7000 to 10,000 cells per sample, following the manufacturer’s guidelines. Subsequently, single-cell RNA-seq libraries were generated utilizing the Chromium Single Cell□3′ v.3.1 kit (10x Genomics, no. PN-1000121). These libraries were sequenced on the NovaSeq□6000 platform (Illumina) to achieve an average of approximately 50,000 reads per cell.

### Bioinformatic analysis

Analysis was performed in the programming languages R (version 4.0.3) and RStudio (version 1.1.463). Data were loaded into R and processed using the Seurat package (version 3.2.3), a comprehensive framework for single-cell omics analysis developed by the Satija Lab. Cells with less than 2500 detected genes were removed, together with cells with more than 15% mitochondrial reads. Filtering using ribosomal genes was not done since these genes did not have a significant effect on clustering (low differences of ribosomal protein-related genes among clusters)

Filtered data were normalized for sequencing depth using the Seurat NormalizeData function with the LogNormalize method and default settings. The variable genes across cells were identified with the Seurat FindVariableFeature function. Differential gene expression between selected groups of cells was assessed using the Seurat FindAllMarkers function with the Wilcoxon Rank Sum test (logfc. threshold = 0.25, min.pct = 0.1, min.diff.pct = 0.2).

### Immunohistochemistry (IHC)

Tumors from patients (see ethics and origin section I.2) were fixed in formalin-fixed, paraffin-embedded (FFPE) blocks and underwent sectioning into 3μm slices using a microtome. Embedded tumor slides were subjected to baking at 60°C for 1 hour and then dewaxed through immersion in sequential xylene and graded ethanol baths using the Jung XL AutoStainer by Leica. Heat-induced epitope retrieval (HIER) was executed in 10 mM citrate buffer (pH = 6) or 10 mM Tris-EDTA buffer (pH = 8) at 125°C for 15 minutes in a pressure cooker.

To inhibit endogenous peroxidase activity, slides were immersed in 0.3% H2O2 for 10 minutes and subsequently blocked in 2.5% w/v BSA in TBS-T for 30 minutes. Primary antibodies (Anti-Netrin from Abcam (AB122903), anti-UNC5B from R&D system (AF1006), in 2.5% w/v BSA in TBS-T) were incubated for 1 hour. Incubation was made with the HRP-conjugated secondary antibody (Anti-Rabbit HRP (MP7401), Anti-Goat HRP (MP7405); in 2.5% w/v BSA in TBS-T) for 30 minutes at room temperature. Chromogenic detection was carried out using the DAB Peroxidase (HRP) Substrate Kit (Vector Laboratories, Burlingame, CA, USA, Ref: SK-4100) following the manufacturer’s instructions.

Following counterstaining with hematoxylin, images were captured using VS120 Virtual Slide Microscope by Olympus. Visualization was accomplished and photographs were taken using the OlyVIA viewer software (v 2.9).

### In Vivo Assays

Five-week-old (15-20 g body weight) female nude mice were obtained from the Charles River RMS Rhone Alpes Auvergne animal facility in France. The mice were housed in sterilised filter-topped cages and maintained in a pathogen-free animal facility. yP190LT organoid cell line was implanted via subcutaneous injection into the right flank of the mice of a solution containing whole organoids in 100 μL of growth factors reduced Matrigel (Corning) diluted in 100 μL of DMEM/F12 medium. Each mouse received a quantity of organoids corresponding to 10 confluent wells in a 48-well plate. To do so, a total of 300 48-wells (for 30 mice) were split and left 72h to reach appropriate confluency, the total amount of organoids was then harvested, washed and resuspended in an appropriate amount of medium/Matrigel to ensure a steady concentration of organoids in each injection. Tumors were measured with a calliper and volume were calculated by the formula V = 0.5 (length × width²). Randomization into groups was established when tumors reached an average of 70 mm3.

Netrin-1 antibody (NP137) or buffer solution used as a control was injected intraperitoneally into mice three times a week at 20 mg/kg once tumors reached 70 mm^3^. The FOLFOX regimen (solution combining 5-Fluoro Uracil (50mg/Kg IP), oxaliplatin (6mg/Kg IP) and disodium levofolinate (90mg/Kg IP)) was injected intraperitoneally, once daily for two days when tumors reached the 70 mm^3^ average. This protocol is routinely used and has been previously reported in the literature by Zoetemelk M. et al^20^ for the treatment of colorectal cancer liver metastases, where it was found to have minimal impact on mice welfare. Tumor volume was monitored at regular intervals (twice a week). After 10 weeks, mice were sacrificed.

### Statistical Analysis

Statistical analyses were performed using GraphPad Prism software (GraphPad, v9.4.1) or R (R, v4.3.2). Data is expressed as the mean, plus or minus SD (Standard Deviation) unless stated otherwise. Data were pooled from a minimum of 3 independent replicated experiments. The threshold for rejecting the null hypothesis was p< 0.05; When statistical significance is integrated into a figure, it is denoted as: * for p< 0.05; ** for p<0.01, *** for p<0.001 and **** for p<0.0001. Paired t-tests were used in *in vitro* experiments with two conditions, cell lines with a different number of passages being considered as a biological replicate. One-way ANOVA, Tukey’s multiple comparison tests was used in *in vitro* experiments with more than two conditions. One-way ANOVA and Tukey’s multiple comparison tests were used to determine the statistical significance difference among mice tumor weight. Two-way ANOVA and Dunnett’s multiple comparison tests were used to determine statistical significance differences among mice tumor growth. The precise details on the type of test used, the number of replicates and the representation of the data are given in the figure’s legends.

## Results

### Netrin-1 and UNC5B profiling in metastatic colorectal cancer

We first assessed the presence of netrin-1 and its receptors in metastatic colorectal cancer (mCRC). Interestingly, analysis of TCGA data showed an increased expression of both netrin-1 and UNC5B mRNA in later stages of CRC, suggesting a potential role for these proteins in metastatic tumors (**Supplementary Figure 1A, 1B**). Immunohistochemistry revealed detectable levels of netrin-1 and of its receptor, UNC5B, in matching mCRC liver metastasis samples (**Supplementary Figure 1C, 1D**). To further explore the role of netrin-1 in mCRC, we conducted bulk RNA sequencing on a cohort of 58 mCRC tumors and 35 paired healthy adjacent liver tissues. The RNA sequencing data revealed a positive correlation between netrin-1 (NTN1) mRNA and UNC5B mRNA expression, suggesting that the dependence receptor/ligand dynamic between these proteins could occur in mCRC (**Figure 1A, 1B**). Moreover, UNC5B emerged as the predominantly expressed netrin-1 receptor in mCRC tumors, showing higher expression compared to other netrin-1 receptors, including DCC, UNC5A, UNC5C, and UNC5D (**Figure 1C**). Interestingly, UNC5B was significantly overexpressed in mCRC tumors compared to the adjacent normal liver tissue (**Figure 1D**), whereas netrin-1 expression was higher in adjacent liver tissue compared to mCRC tumors (**Figure 1E**), implying that the liver could serve as a local microenvironmental source of netrin-1 for tumor cells in mCRC liver metastases, in line with what was reported recently for metastatic pancreatic cancer^21^.

**Figure 1.**
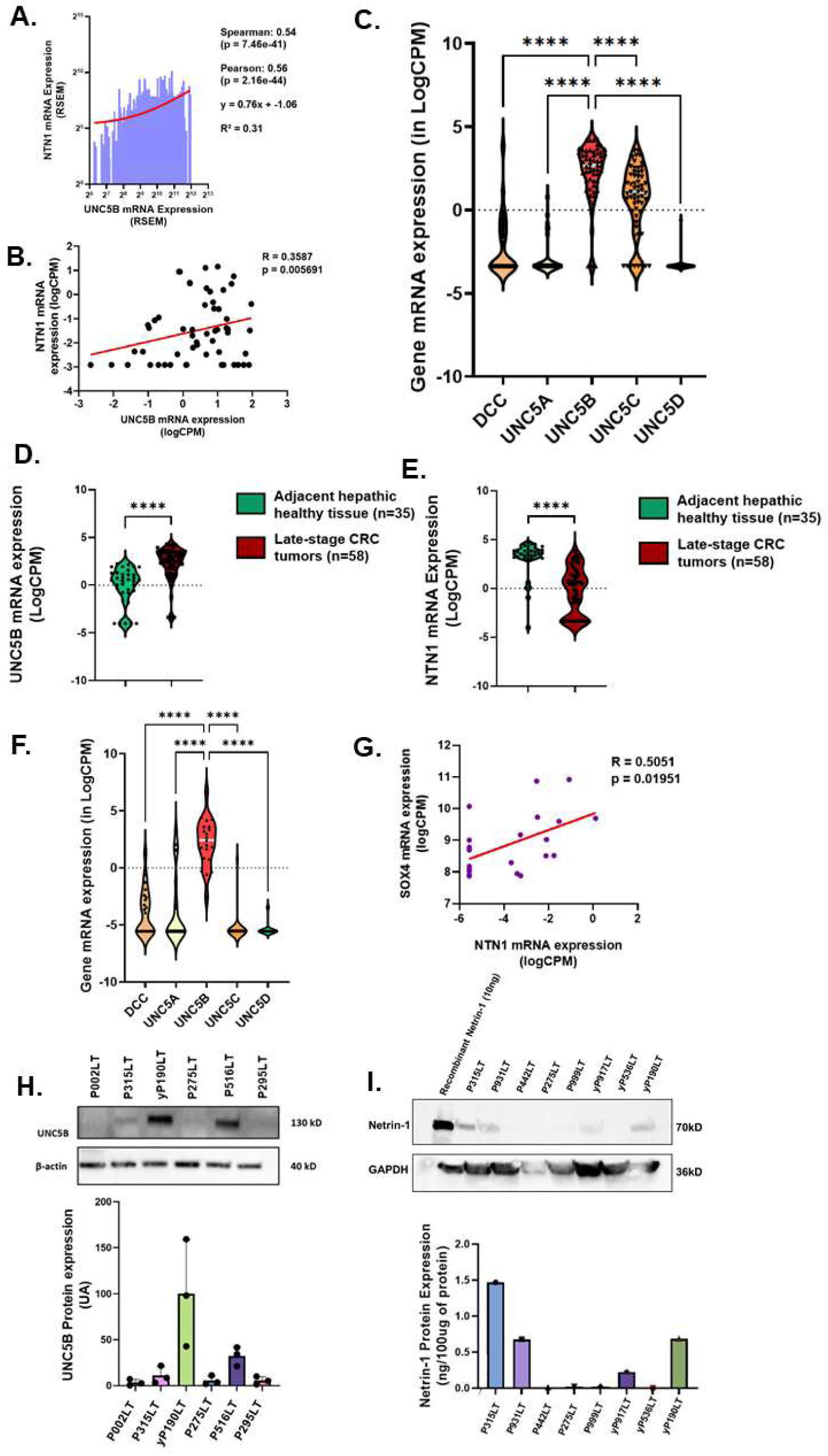
Comparative Analysis of netrin-1 (NTN1) and UNC5B expression in stage IV CRC tumors and PDOs -. **(A)** Correlation between UNC5B and NTN1 mRNA expression in the TCGA colon adenocarcinoma dataset; **(B)** Correlation between UNC5B mRNA and NTN1 mRNA expression in stage IV CRC tumors; (**C**) Expression of netrin-1 receptors (DCC, UNC5A-D) mRNA in stage IV CRC tumors (n=58) (bars indicate means); **(D-E)** Comparison of mRNA Expression in Healthy adjacent liver and Stage IV colorectal tumor tissues for UNC5B **(D)** and NTN1 **(E)**; **(F)** Summary of mRNA expression levels of various netrin-1 receptors in organoids derived from metastatic colorectal tumors (n=21); **(G)** Correlation between UNC5B mRNA and NANOG mRNA expression; **(H-I)** Quantification of UNC5B **(H)** and netrin-1 **(I)** protein expression in organoid cell lines using Western blotting; Top panels: representative Western Blot image; Bottom panel: quantification of protein expression across organoid cell lines; Normalized gene expression is expressed in logCPM; (TCGA cohort: n=521; stage IV tumors cohort: n=58, Adjacent healthy tissue cohort: n=34); p Values: * = <0.05, ** = <0.01, **** = <0.0001; Statistical test: Spearman/Pearson for (A); Spearman’s rank correlation coefficient for (B); One-Way ANOVA - Tukey’s multiple comparisons test for (C) and (F), unpaired t-tests for (D) and (E); (H) Results are expressed as a % of yP190LT mean expression, using β-actin as loading control; 3 replicates per organoid cell line. (I) Results are expressed as % compared to lane containing 20ng of Recombinant netrin-1, using GAPDH as loading control; 1 replicate per organoid cell line.

Aiming to develop a reliable preclinical model to study the role of netrin-1/UNC5B, we analyzed their expression in patient-derived organoids (PDOs) derived from surgical liver metastasis specimens of stage IV CRC patients. RNA sequencing confirmed the expression of UNC5B, as the predominant receptor, in the 21 PDOs analyzed (**Figure 1F**), consistent with what we observed in mCRC tumors. Moreover, we identified a positive correlation between UNC5B and the stemness marker SOX4 at the mRNA level (**Figure 1G**). We further confirmed the expression of netrin-1 and UNC5B proteins in a subset of PDOs using Western blot analyses (**Figure 1H, 1I**).

### Netrin-1 and UNC5B modulates mCRC cells self-renewal

Based on the view that netrin-1 may act in a paracrine manner in liver mCRC, we assessed the effect of recombinant netrin-1 on the self-renewal capacity of a set of PDOs expressing UNC5B (yP190LT, P315LT, and P931LT) using extreme limiting dilution assays (ELDAs)(**Figure 2A, 2B, 2C**)^19^. Netrin-1 (100ng/mL) significantly enhanced self-renewal, a core feature of CSCs, across all three PDOs. This effect was fully reversed by the netrin-1 neutralizing antibody NP137 (10ug/mL), supporting the role of netrin-1 in promoting self-renewal. Of interest, NP137 alone significantly decreased self-renewal compared to both untreated and control antibody (NP001)-treated conditions in netrin-1 expressing CRC cell lines such as HCT116 (**Supplementary Figure 2**), supporting the view that endogenous netrin-1 promotes self-renewal.

**Figure 2.**
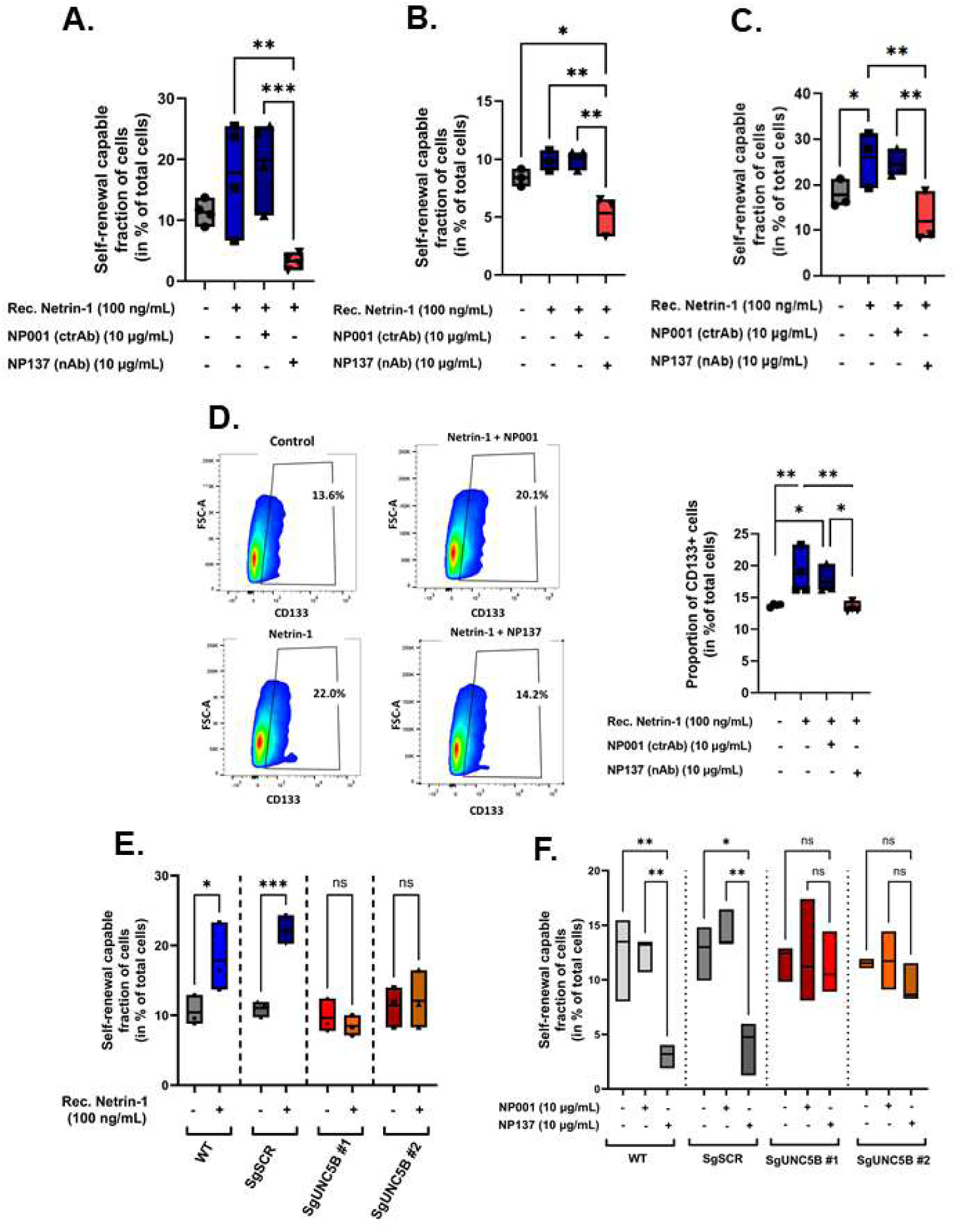
Netrin-1 enhances self-renewal in mCRC organoids through the activation of UNC5B -. **(A-C)** Self-renewal capable fraction (as a % of total cell number) in yP190LT **(A),** P315LT **(B)** and P931LT **(C)** organoids in the presence or absence of recombinant netrin-1 (100 ng/mL), NP001 (10μg/mL) and/or NP137 (10μg/mL), quantified using ELDA; (**D**) (Left) Representative flow cytometry analysis (Gate was determined in comparison with an unstained control and left unchanged for all samples) and (Right) bar graph summarizing proportions of CD133-positive cells in yP190LT organoid after 72h of treatment with recombinant netrin-1 (100 ng/mL), NP001 (10μg/mL) and/or NP137 (10μg/mL); **(E)** Self-renewal capable fraction of in wildtype, scramble control (sgSCR) and CRISPR-edited (SgUNC5B#1 and SgUNC5B#2) P315LT organoids, either left untreated (Control) or treated with recombinant netrin-1 (100 ng/mL), observed by ELDA; **(F)** Self-renewal capable fraction of in wildtype, scramble control (sgSCR) and UNC5B-null (SgUNC5B#1 and SgUNC5B#2) P315LT organoid cells, untreated (Control) or treated with NP001 (10μg/mL) or NP137 (10μg/mL), quantified using ELDA; Statistical significance was determined using One-way Anova – Tukey’s multiple comparison test, ns = non-significant, * = p <0.05, ** = p <0.01, *** = p <0.001. N=3 for (B), (C), (E) and (F)); N=4 for (A) and (D)

We further explored the effect of netrin-1 on the CSCs population by quantifying the proportion of cells expressing CD133, a marker often associated with CSCs in colorectal cancer, in organoids expressing the highest level of UNC5B (yP190LT). We observed that netrin-1 supplementation significantly increased the proportion of CD133-positive cells, an effect that was inhibited by NP137 (**Figure 2D**).

To demonstrate the importance of UNC5B in the regulation of self-renewal by netrin-1, we next silenced UNC5B using CRISPR-Cas9 technology in P315LT PDOs (**Supplementary Figure 3**). We observed that UNC5B silencing did not significantly affect self-renewal in the absence of exogenous netrin-1. However, the promotion of self-renewal by recombinant netrin-1 was completely abolished in UNC5B-null PDOs (SgUNC5B), in comparison with wild-type and control (SgSCRAMBLE) PDOs (**Figure 2E**). This data demonstrates that UNC5B is essential to mediate the effect of netrin-1 on self-renewal. Additionally, the inhibitory effect of NP137 in wild-type and SgSCRAMBLE PDOs was abolished in SgUNC5B PDOs (**Figure 2F**), indicating that this effect relies on the dependence receptor activity of UNC5B. Together these data support the view that, in mCRC liver metastases, self-renewal is promoted by netrin-1 via an UNC5B dependent mechanism.

### Netrin-1 promotes Self-Renewal Through Apoptosis regulation

Next, we further investigated how netrin-1 influences self-renewal via UNC5B. UNC5B has been shown to display at least two opposing signaling activities depending on netrin-1 presence/absence, including the activation of cell death in various settings when its ligand is absent^22–24^. We thus looked more specifically at apoptosis induction in CD133+ cells of PDOs treated with netrin-1 and/or with the netrin-1 mAb. Flow cytometry analysis of Annexin V – Propidium Iodide staining highlighted a significant decrease in late-apoptosis among CD133+ cells treated with netrin-1 (**Figure 3A**). This anti-apoptotic effect was fully reversed by the addition of NP137, supporting the view that netrin-1 promotes CSC survival. Of note, no such protective effect of netrin-1 on apoptosis was observed in CD133-negative cells (**Figure 3B**).

**Figure 3.**
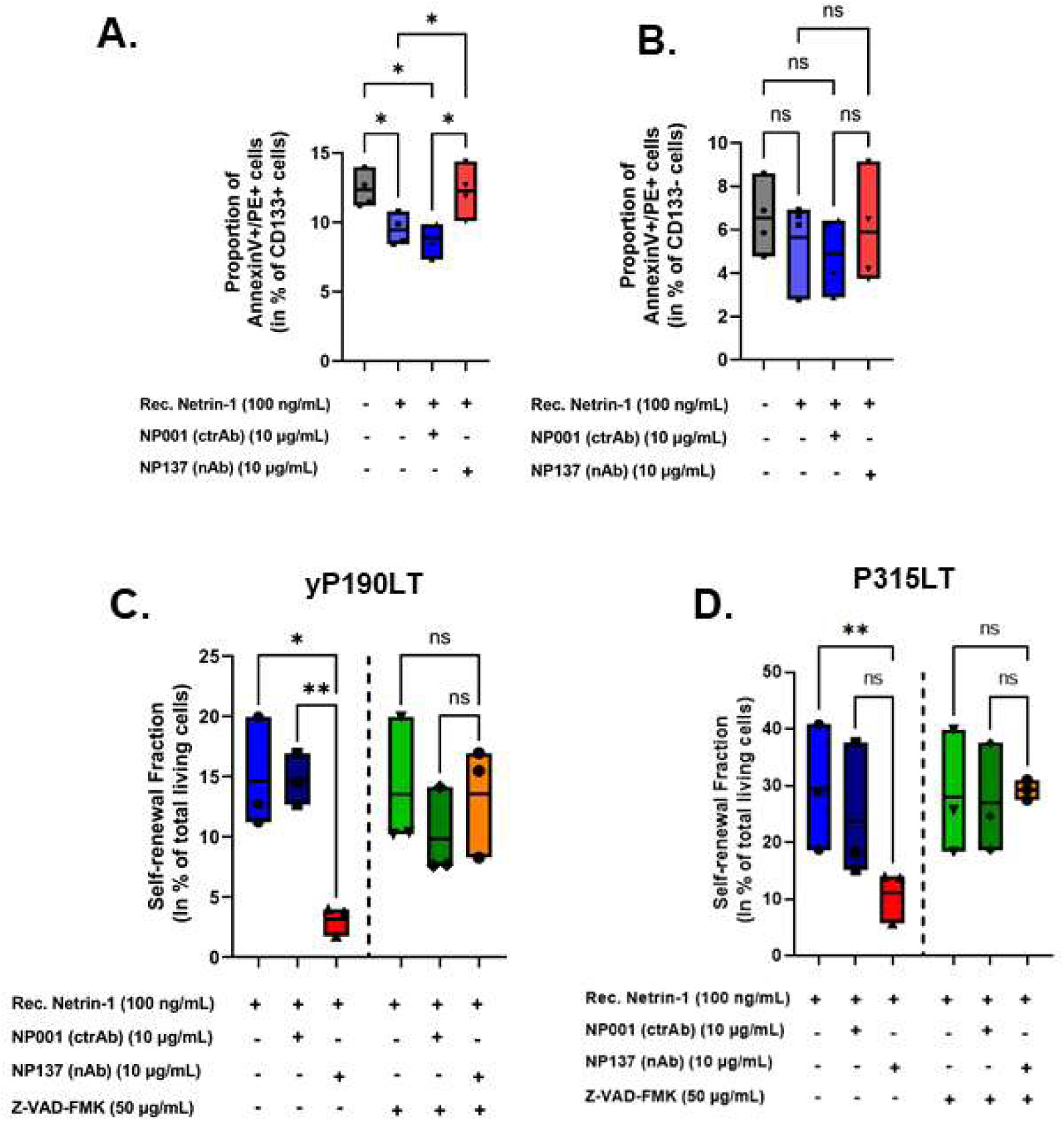
Netrin-1 modulates self-renewal in mCRC through CSC apoptosis -. **(A)** Flow cytometry quantification of late apoptosis in CD133-positive cells from yP190LT organoid after 72h of treatment with recombinant netrin-1 (100 ng/mL), NP001 (10μg/mL) and/or NP137 (10μg/mL) as determined by annexing V / Propidium iodide double staining; **(B)** Flow cytometry quantification of late apoptosis in CD133-negative cells from yP190LT organoid after 72h of treatment with recombinant netrin-1 (100 ng/mL), NP001 (10μg/mL) and/or NP137 (10μg/mL) as determined by annexing V / Propidium iodide double staining; **(C-D)** Self-renewal capable fraction of yP190LT**(C)** and P315LT**(D)** organoid cell line in presence or absence of recombinant netrin-1 (100 ng/mL), NP001 (10μg/mL) and/or NP137 (10μg/mL) and Z-VAD-FMK (50ug/mL) quantified using ELDA; Statistical significance was determined using One-way Anova – Tukey’s multiple comparison test, Error bars indicate SD; ns = non-significant, * = p <0.05, ** = p <0.01, **** = p <0.0001. N=4 for (a) and (b), N=3 for (c)

The essential role of apoptosis regulation in netrin-1-mediated self-renewal promotion was further corroborated using a caspase inhibitor, z-VAD-FMK. In both yP190LT and P315LT organoids, the caspase inhibitor fully blocked the effect of NP137 on self-renewal, demonstrating that the main mode of action of this therapeutic netrin-1 blocking antibody is to induce CSC apoptosis (**Figure 3C, 3D**).

### TFF3 acts as paracrine factor to promote netrin-1 mediated self-renewal

To further understand the mechanism underpinning netrin-1/UNC5B role on self-renewal and CSC survival, we performed scRNAseq on yP190LT PDOs treated with NP137 or with an isotype mAb control. In line with our previous observations, a significant decrease in the expression of key stemness-related genes such as LGR5, SMOC2, SOX4, and PROM1 was observed in NP137-treated organoids (**Figure 4A-B**). Gene ontology analysis indicated that pathways associated with cell adhesion and development were impaired by netrin-1 blockade (**Figure 4C**). Unexpectedly, cancer cells enriched for UNC5B receptor expression were largely different from those exhibiting high levels of stemness markers such as LGR5 (**Figure 5A**). Moreover, the decrease in stemness-related genes following NP137 treatment was not seen specifically in UNC5B-expressing cells (**Figure 5B**), suggesting that while NP137 blocks the binding of netrin-1 and at the surface of UNC5B expressing cells, the full impact of its activity on CSC survival and self-renewal may require a paracrine mechanism.

**Figure 4.**
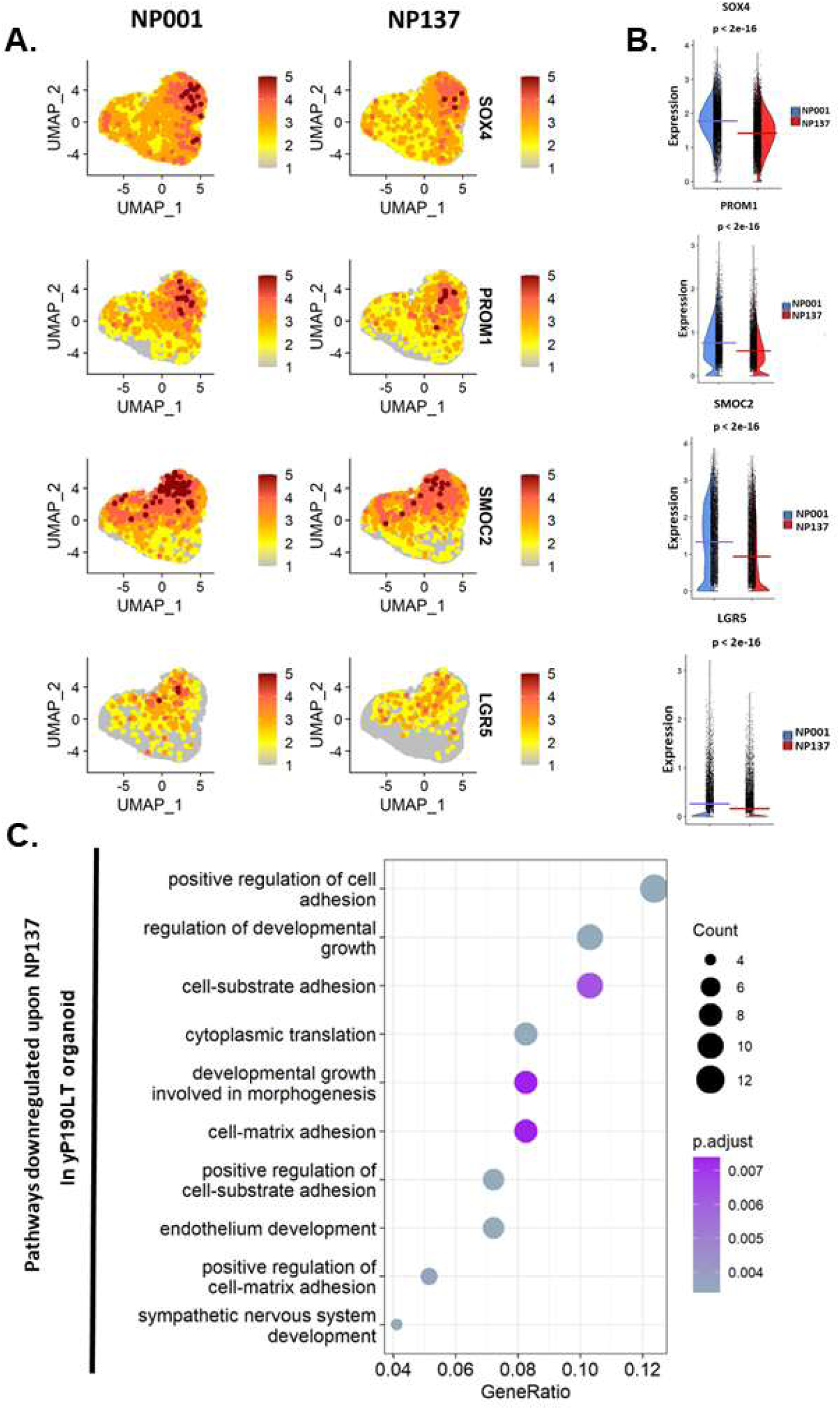
NP137 impairs expression of stemness-related genes in yP190LT – scRNA sequencing analysis -. **(A)** UMAP plot summarizing the expression of SOX4, SMOC2, PROM1 and LGR5 in yP190LT organoids after 72h of treatment with NP001 (10μg/mL) or NP137 (10μg/mL) (Means and p.values displayed determined by stat_compare_means function; Bars indicate means, statistical test use : Wilcoxon test, precise values displayed in table 5-1); **(B)** Violin plot of SOX4, SMOC2, PROM1 and LGR5 expression level in yP190LT organoids after 72h of treatment with NP001 (10μg/mL) or NP137 (10μg/mL); (**C**) Pathway enrichment analysis for genes downregulated in yP190LT upon NP137 treatment (Genes differentially expressed determined using FindMarkers function of Seurat package, pathways determined using enrichGO function of clusterProfiler package, Gene Ratio is a metric used to measure the degree of enrichment of a gene set within a larger gene list. It is calculated by dividing the number of genes from the gene set that are present in the input gene list by the total number of genes in the gene set; Genes were considered differentially expressed if log2foldchange>0.15; p.value cut. off = 0.04; Statistical test used: Benjamini-Hochberg) Means and p.values displayed determined by stat_compare_means function; Bars indicate means, statistical test use : Wilcoxon test, each dot represent an event (cell))

**Figure 5.**
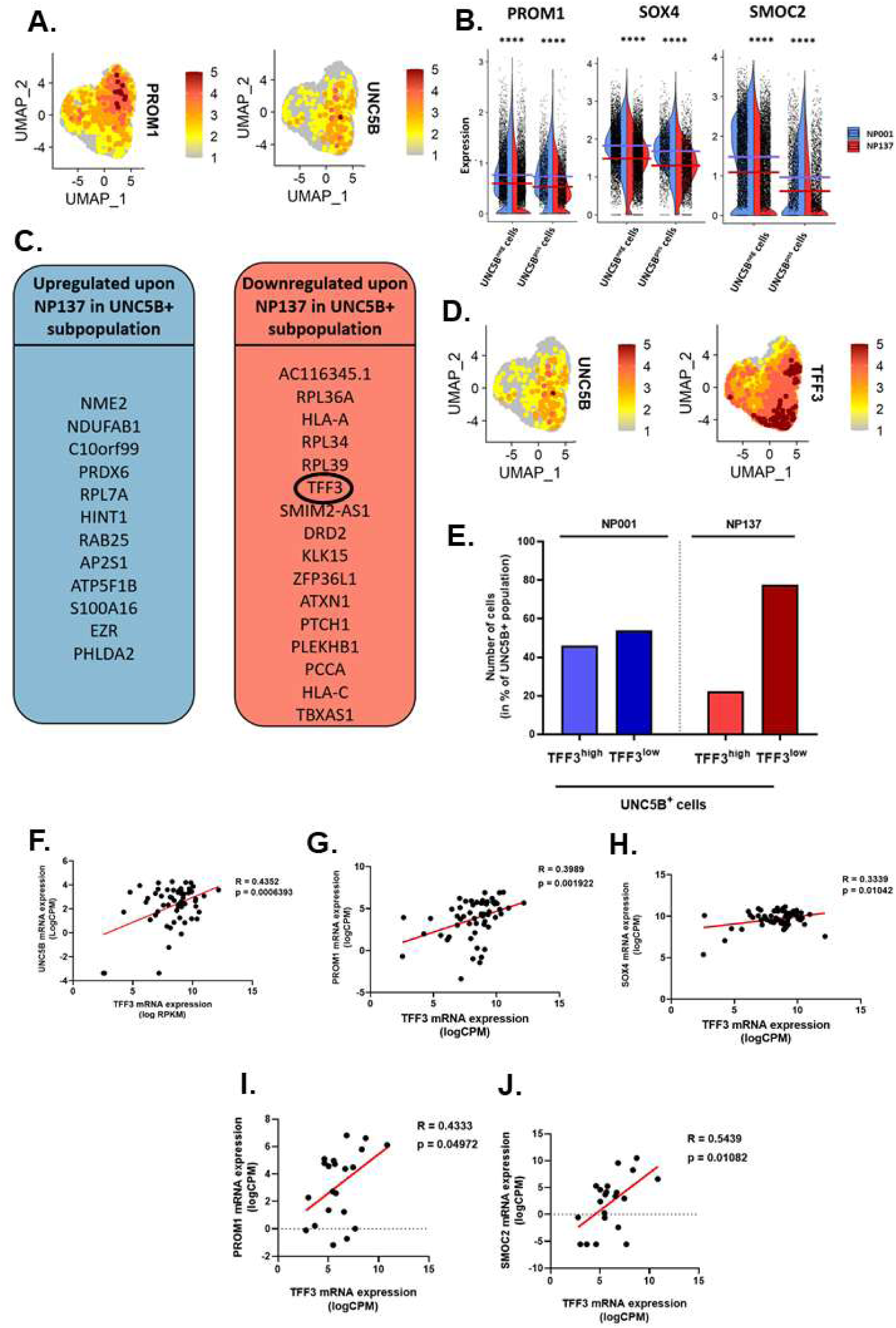
TFF3 expression is decreased by NP137 treatment and correlates with stemness-related genes -. **(A)** UMAP plot of **LGR5** and **UNC5B** expression level in control yP190LT organoid cells (NP001-treated); **(B)** Violin plot of the expression of **PROM1, SOX4 and SMOC2 genes** in UNC5B^neg^ and UNC5B^pos^ cells treated with either NP001 or NP137 (Each dot represent an event (cell)), **(C)** Fraction of high-expressing TFF3 and low-expressing TFF3 in UNC5B-positive cells treated with either NP001 (10μg/mL) or NP137 (10μg/mL); **(D)** UMAP plot of **UNC5B** (Left) and **TFF3** (Right) expression level yP190LT organoid cells (treated with NP001 control antibody (10μg/mL)); **(E)** UMAP plot of **TFF3** expression level yP190LT organoid cells treated with either NP001 or NP137; **(F)** Correlation between UNC5B mRNA and **TFF3** in stage IV CRC tumors **(G)** Correlation between PROM1 mRNA and TFF3 in stage IV CRC tumors; **(H)** Correlation between SOX4 mRNA and TFF3 in stage IV CRC tumors; **(I)** Correlation between PROM1 mRNA and TFF3 mRNA in mCRC organoids; **(J)** Correlation between SMOC2 mRNA and TFF3 mRNA in mCRC organoids; For (C) High TFF3 expression is defined as > log5 and low expression as < log4; Means and p.values displayed determined by stat_compare_means function; Bars indicate means, statistical test use: Wilcoxon test for (C) and (E), Spearman’s rank correlation coefficient for (G-I); One-way Anova – Tukey’s multiple comparison test for (k), *** = p<0.001, **** = p<0.0001

We thus mined our scRNAseq data for genes that may be specifically decreased in UNC5B-expressing cells upon treatment with NP137 (**Figure 5C**). The only extracellular communication related gene significantly impacted was Trefoil Factor 3 (TFF3), a secreted peptide involved in mucosal protection and repair and known for its role in promoting stemness in cancer cells ^17,25–28^. Elevated TFF3 levels have been associated with increased stemness, resistance to apoptosis, and enhanced tumorigenic potential in various cancers^16,25–28^. Importantly, high baseline levels of TFF3 were detected in cells expressing the UNC5B receptor (**Figure 5D**) and NP137 treatment selectively induced a decrease of the fraction of cells expressing high levels of TFF3 within the UNC5B-positive cell subset (**Figure 5E**). Whereas no effect of NP137 treatment on the fraction of cells expressing high levels of TFF3 within the UNC5B-negative cell subset was observed (**Supplementary Figure 4**).

Analysis of bulk RNA sequencing data from the cohort of mCRC tumors and organoids revealed a positive correlation between TFF3 mRNA and UNC5B mRNA expression (**Figure 5F**). Supporting an association between TFF3 and stemness in mCRC, a strong correlation was observed between the mRNA expression of TFF3 and several stemness-related genes such as PROM1, SOX4 and SMOC2 in PDOs and tumor samples (**Figure 5G – 5J**). Together, this suggests that TFF3 may act downstream of netrin-1/UNC5B signaling to regulate stemness in mCRC cells.

To explore this formally we moved back to PDOs culture. Using ELISA assays, we observed that recombinant netrin-1 significantly increased the secretion of TFF3 in PDOs, including yP190LT, P315LT, and P931LT (**Figure 6A**). This effect is independent from PDO proliferation (**Supplementary Figure 5A**) and requires netrin-1 binding to the UNC5B receptor, as PDOs lacking UNC5B did not show any increase in TFF3 secretion upon netrin-1 stimulation (**Supplementary Figure 5B).** To further explore whether TFF3 is involved as a mediator of netrin-1 activity on self-renewal, we blocked its activity using AMPC, an inhibitor of TFF3 dimerization^17^. Inhibition of TFF3 activity led to a significant decrease in self-renewal capacity in the PDOs, reaching levels comparable to those observed in samples treated with the NP137 antibody (**Figure 6B**). Critically, netrin-1 was no longer able to enhance self-renewal in the presence of AMPC and combining NP137 with AMPC did not provide any additive or synergistic inhibitory effect, altogether suggesting that TFF3 is a critical downstream effector of self-renewal modulation by netrin-1/UNC5B.

**Figure 6.**
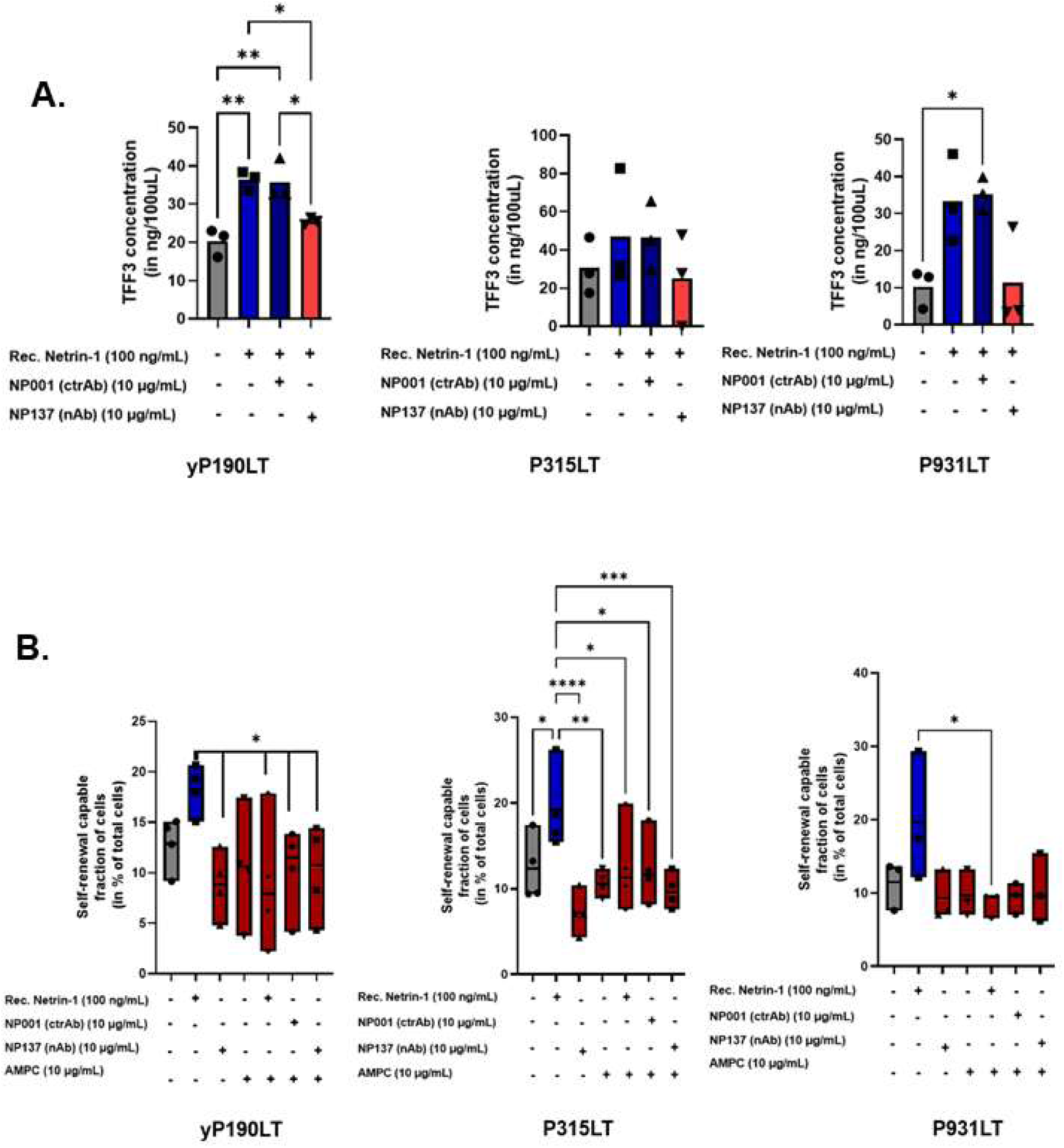
Netrin-1 enhances self-renewal though TFF3 -. **(A)** TFF3 expression in the conditioned medium from liver metastasis PDOs (yP190LT; P315LT; P931LT) treated or not with recombinant netrin-1 (100 ng/mL), NP001 (10μg/mL) and/or NP137 (10μg/mL), quantified using ELISA (n=3); **(B)** Self-renewal capable fraction of P931LT (Left) and P315LT (Right) organoid cell line in presence or absence of recombinant Netrin-1 (100 ng/mL), NP001 (10μg/mL), NP137 (10μg/mL) and/or AMPC (10ug/mL), quantified using ELDA; Statistical significance was determined using One-way Anova – Tukey’s multiple comparison test, Error bars indicate SD; ns = non-significant, * = p <0.05, ** = p <0.01, **** = p <0.0001. N=3

Of interest, NP137 was recently used as a single agent to induce netrin-1 blockade in patients with advanced cancers in a phase 1 clinical trial^15^. From 42 patients included in this trial, only 2 had colorectal cancer and only one had liver metastasis expressing detectable levels of NTN1 and TFF3 genes. RNAseq was performed on biopsies collected both prior to (baseline, C1D1) and after a 1-month systemic treatment with two doses of NP137 (C3D1) (**Figure 7A**). Similarly, to what was observed in PDOs, the high level of TFF3 detected at baseline was dramatically reduced after NP137 treatment, together with that of multiple stemness-related genes, including SOX4, PROM1, SMOC2, LGR5, and EPCAM (**Figure 7B**). Together, these results indicate that the netrin-1/UNC5B axis promotes self-renewal in mCRC cells through the secretion of TFF3.

**Figure 7.**
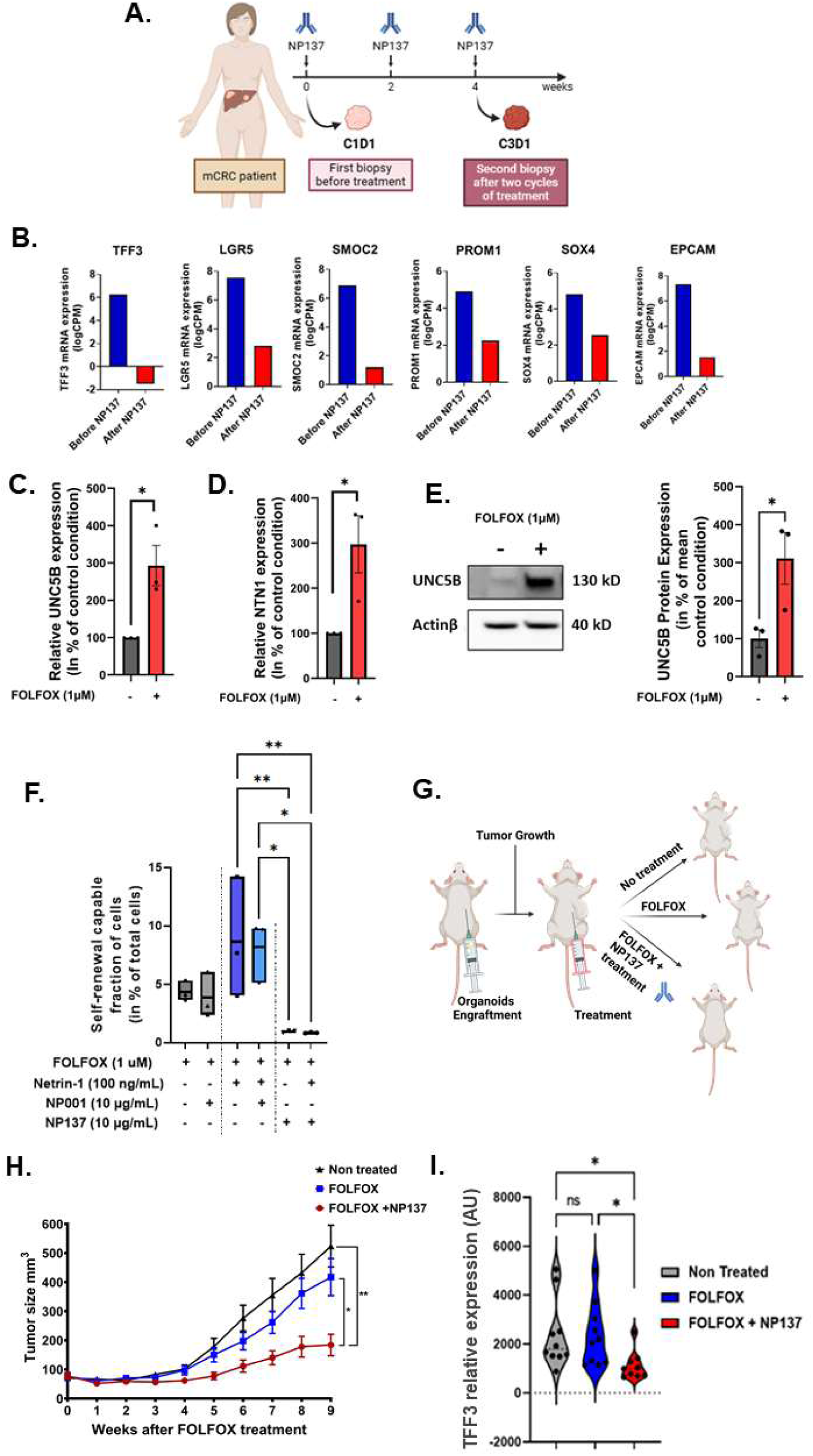
Targeting Netrin-1 to enhance FOLFOX treatment in mCRC -. **(A)** Diagram summarizing the longitudinal collection protocol of liver metastasis biopsies prior to (C1D1) or after (C3D1) systemic treatment with NP137 in a patient with mCRC; **(B)** TFF3, PROM1, LGR5, SMOC2, SOX4 and EPCAM mRNA in CRC tumor of a patient treated with NP137 levels determined by RNA sequencing; Data analyzed using DESseq2 package in R; **(C)** Quantification of relative **UNC5B** mRNA expression level in yP190LT PDO after 24h of treatment with either the vehicle or FOLFOX (1 µM) by real-time qPCR; **(D)** Real-time qPCR quantification of relative **NTN1** mRNA expression level in yP190LT PDO after 24h of treatment with either vehicle or FOLFOX (1 µM); **(E)** Western blot quantification of **UNC5B** protein expression level in yP190LT PDO after 72h of treatment with either vehicle or FOLFOX (1 µM) followed by a 15-day recovery period; **(F)** Self-renewal capable fraction of yP190LT organoid cell line after 72h of FOLFOX treatment (72h; 1 µM) in presence or absence of recombinant Netrin-1 (100 ng/mL), NP001 (10μg/mL) and/or NP137 (10μg/mL) observed by ELDA; Statistical test used : One-way Anova – Tukey’s multiple comparison test **(G);** Schematic representation of In Vivo protocol to evaluate the therapeutical potential of NP137 in mCRC using PDO-xenografted mice; **(H)** Evolution of tumor size in BALBc nude mice subcutaneously engrafted with yP190LT PDO after treatment with either PBS or FOLFOX (solution combining 5-Fluoro Uracil (50mg/Kg IP), oxaliplatin (6mg/Kg IP) and disodium levofolinate (90mg/Kg IP) daily for two days) or FOLFOX and NP137 (20 mg/kg; three times a week); N = 8/group, Error bars indicate SEM; Statistical test used : Two-way Anova - Dunnett’s multiple comparisons test; **(I)** Real-time qPCR quantification of TFF3 expression of engrafted tumors in mice treated with either PBS, FOLFOX or FOLFOX and NP137 at week 10; N = 8/group, Error bars indicate SD, Statistical test used : One-way Anova - Dunnett’s multiple comparisons test

### Targeting Netrin-1 to Enhance FOLFOX Treatment in mCRC

Based on the known role of cancer stemness in chemoresistance, we next investigated the potential of targeting netrin-1 in combination with FOLFOX chemotherapy, a standard of care in mCRC. First, we observed that FOLFOX treatment induced a notable increase in netrin-1 and UNC5B mRNA expression (**Figure 7C, 7D**). Increased expression of UNC5B was maintained for at least 20 days after FOLFOX treatment, as detected using Western blotting (**Figure 7E**). Given these observations, we hypothesized that inhibiting netrin-1 could improve the effect of FOLFOX by further reducing self-renewal capacity in PDOs. Indeed, we observed a significant downregulation of self-renewal in PDOs treated with NP137 in combination with FOLFOX, compared to those treated with either recombinant netrin-1 and the control antibody NP001 (**Figure 7F**). This finding suggests that NP137 could improve the effect of chemotherapy by targeting a subset of self-renewing cells that usually survive FOLFOX exposure.

To evaluate the pre-clinical relevance of this combination treatment, we engrafted the yP190LT PDO into BALB/c nude mice and subjected them to three treatment conditions: no treatment, FOLFOX alone, and FOLFOX combined with NP137 (**Figure 7G**). We observed a significant reduction in tumor growth in the FOLFOX + NP137 treatment group compared to both untreated and FOLFOX-only treated mice (**Figure 7H, Supplementary Figure 6B**). Tumors from the NP137-treated group also had significantly lower weight compared to those from the other two treatment groups (**Supplementary Figure 6A, 6B**). Interestingly, in tumours treated with NP137, TFF3 expression was significantly reduced at week 10 compared to those treated with PBS or FOLFOX alone (**Figure 7I**), supporting the hypothesis that netrin-1/UNC5B also control TFF3 expression within mCRC tumours *in vivo*. Collectively these results highlight the potential of targeting the netrin-1/UNC5B interaction to improve the efficacy of currently used chemotherapy like FOLFOX in mCRC, providing a promising therapeutic strategy to overcome resistance mechanisms associated with cancer stem cells and tumor recurrence.

## Discussion

In this study, we have demonstrated the key role of netrin-1/UNC5B in the self-renewal, stemness, and chemoresistance of mCRC cells using a series of expression dataset and experimentations in patient derived organoids. The findings build on previous works that highlighted the involvement of netrin-1 in tumorigenesis and epithelial-mesenchymal transition in various solid tumors. By extending these insights to mCRC, our data provide a comprehensive view of how netrin-1 influences critical aspects of cancer cell behavior and suggests that targeting netrin-1 could be an effective therapeutic strategy especially in combination with chemotherapy.

How netrin-1 regulates stemness and plasticity has so far not been addressed. Of interest we succeeded in showing that netrin-1 acts as a survival factor for CD133+ cancer cells in mCRC. This is also the first demonstration of the mode of action of the therapeutic antibody which is, by disrupting netrin-1/UNC5B interaction^29^, inducing more specifically cell death in CD133+ cells. However, in a rather unexpected manner it does not appear to be a direct effect. Indeed, we provide a series of compelling data showing that at the single cell level, the cancer cells which are impacted by the NP137 treatment are not the one expressing UNC5B. We uncover a paracrine mechanism where UNC5B expressing cancer cells secrete TFF3 upon netrin-1 binding resulting in CSCs survival. Not only this provides new insights in a mechanism that was though direct^30^, but it also potentially provides a potential biomarker to stratify patients that could respond to NP137. Indeed, the NP137 is currently assessed in a variety of phase 1b/2 clinical trials evaluating the netrin-1 blockade in combination with chemotherapies and/or immune checkpoint (clinical.gov NCT04652076; NCT05605496; NCT05546853; NCT05546879), however based on the data shown here, it would worth assessing netrin-1 blockade in combination with FOLFOX specifically in patients showing high level of TFF3. The patients showing no TFF3 or low level of TFF3 should not benefit of the NP137 treatment as shown here with the PDOs treated with the TFF3 inhibitor. On the contrary, the patients showing high level of TFF3 should be the ones who will benefit most of netrin-1 blockade as it should reduce TFF3 level, leading to reduced CSCs survival and thus less resistance to FOLFOX. Whether this is restricted to liver metastasis of mCRC or a general mechanism for netrin-1/UNC5B function in cancer plasticity remains to be further investigated.

## Data Availability Statement

All data supporting the findings of this study are available within the paper and its Supplementary data. The TCGA datasets were derived from sources in the public domain and are available on cBioportal website (https://www.cbioportal.org/study/summary?id=coadread_tcga_pan_can_atlas_2018).

## Acknowledgement

We wish to thank the core facilities of CRCL. We also extend our thanks to the Peter MacCallum Cancer Centre for their support through the CAHM (Centre for Advanced Histology and Microscopy) and the Molecular Genomics Core. We acknowledge the Victorian Centre for Functional Genomics for their contributions.

## Authors Contribution

Investigation, M.B, K.R, A.P, L.S, R.W, C.D, F.L, L.F, A.G., C.B and T.V; Conceptualization, M.B., P.M. and F.H.; Resources, C.D., A.H., C.B and T.V; Supervision, P.M. and F.H.; Writing – Original Draft, M.B., P.M. and F.H.; Writing – Review & Editing, M.B., P.M. and F.H.; Funding Acquisition, P.M and F.H.

## References

1. Bray F, Ferlay J, Soerjomataram I, Siegel RL, Torre LA, Jemal A. Global cancer statistics 2018: GLOBOCAN estimates of incidence and mortality worldwide for 36 cancers in 185 countries. CA: A Cancer Journal for Clinicians. 2018;68(6):394–424. 10.3322/caac.21492

2. Rawla P, Sunkara T, Barsouk A. Epidemiology of colorectal cancer: incidence, mortality, survival, and risk factors. Prz Gastroenterol. 2019;14(2):89–103. doi:10.5114/pg.2018.81072

3. Siegel RL, Miller KD, Jemal A. Cancer statistics, 2016. CA: a cancer journal for clinicians. 2016;66(1):7–30.

4. Stintzing S. Management of colorectal cancer. F1000Prime Rep. 2014;6:108. doi:10.12703/p6-108

5. Fakih MG. Metastatic Colorectal Cancer: Current State and Future Directions. Journal of Clinical Oncology. 2015;33(16):1809–1824. doi:10.1200/jco.2014.59.7633

6. Brisset M, Mehlen P, Meurette O, Hollande F. Notch receptor/ligand diversity: contribution to colorectal cancer stem cell heterogeneity. Review. Frontiers in Cell and Developmental Biology. 2023-October-04 2023;11 doi:10.3389/fcell.2023.1231416

7. He Y-C, Zhou F-L, Shen Y, Liao D-F, Cao D. Apoptotic death of cancer stem cells for cancer therapy. International journal of molecular sciences. 2014;15(5):8335–8351.

8. Colak S, Zimberlin C, Fessler E, et al. Decreased mitochondrial priming determines chemoresistance of colon cancer stem cells. Cell Death & Differentiation. 2014;21(7):1170–1177.

9. Serafini T, Kennedy TE, Galko MJ, Mirzayan C, Jessell TM, Tessier-Lavigne M. The netrins define a family of axon outgrowth-promoting proteins homologous to C. elegans UNC-6. Cell. Aug 12 1994;78(3):409–24. doi:10.1016/0092-8674(94)90420-0

10. Deiner MS, Kennedy TE, Fazeli A, Serafini T, Tessier-Lavigne M, Sretavan DW. Netrin-1 and DCC Mediate Axon Guidance Locally at the Optic Disc: Loss of Function Leads to Optic Nerve Hypoplasia. Neuron. 1997/09/01/ 1997;19(3):575–589. 10.1016/S0896-6273(00)80373-6

11. Llambi F, Causeret F, Bloch-Gallego E, Mehlen P. Netrin-1 acts as a survival factor via its receptors UNC5H and DCC. Embo j. Jun 1 2001;20(11):2715–22. doi:10.1093/emboj/20.11.2715

12. Bernet A, Fitamant J. Netrin-1 and its receptors in tumour growth promotion. Expert Opin Ther Targets. Aug 2008;12(8):995–1007. doi:10.1517/14728222.12.8.995

13. Brisset M, Grandin M, Bernet A, Mehlen P, Hollande F. Dependence receptors: new targets for cancer therapy. EMBO Mol Med. 2021;13(11):e14495.

14. Lengrand J, Pastushenko I, Vanuytven S, et al. Pharmacological targeting of netrin-1 inhibits EMT in cancer. Nature. 2023/08/01 2023;620(7973):402–408. doi:10.1038/s41586-023-06372-2

15. Cassier PA, Navaridas R, Bellina M, et al. Netrin-1 blockade inhibits tumour growth and EMT features in endometrial cancer. Nature. 2023/08/01 2023;620(7973):409–416. doi:10.1038/s41586-023-06367-z

16. You ML, Chen YJ, Chong QY, et al. Trefoil factor 3 mediation of oncogenicity and chemoresistance in hepatocellular carcinoma is AKT-BCL-2 dependent. Oncotarget. Jun 13 2017;8(24):39323–39344. doi:10.18632/oncotarget.16950

17. Guo H, Tan YQ, Huang X, et al. Small molecule inhibition of TFF3 overcomes tamoxifen resistance and enhances taxane efficacy in ER+ mammary carcinoma. Cancer Lett. Nov 28 2023;579:216443. doi:10.1016/j.canlet.2023.216443

18. Doench JG, Fusi N, Sullender M, et al. Optimized sgRNA design to maximize activity and minimize off-target effects of CRISPR-Cas9. Nat Biotechnol. Feb 2016;34(2):184–191. doi:10.1038/nbt.3437

19. Hu Y, Smyth GK. ELDA: extreme limiting dilution analysis for comparing depleted and enriched populations in stem cell and other assays. J Immunol Methods. Aug 15 2009;347(1-2):70–8. doi:10.1016/j.jim.2009.06.008

20. Zoetemelk M, Ramzy GM, Rausch M, et al. Optimized low-dose combinatorial drug treatment boosts selectivity and efficacy of colorectal carcinoma treatment. Mol Oncol. Nov 2020;14(11):2894–2919. doi:10.1002/1878-0261.12797

21. Dudgeon C, Casabianca A, Harris C, et al. Netrin-1 feedforward mechanism promotes pancreatic cancer liver metastasis via hepatic stellate cell activation, retinoid, and ELF3 signaling. Cell Reports. 2023;42(11)

22. Guenebeaud C, Goldschneider D, Castets M, et al. The dependence receptor UNC5H2/B triggers apoptosis via PP2A-mediated dephosphorylation of DAP kinase. Mol Cell. Dec 22 2010;40(6):863–76. doi:10.1016/j.molcel.2010.11.021

23. Akino T, Han X, Nakayama H, et al. Netrin-1 Promotes Medulloblastoma Cell Invasiveness and Angiogenesis, and Demonstrates Elevated Expression in Tumor Tissue and Urine of Patients with Pediatric Medulloblastoma. Cancer Research. 2014;74(14):3716–3726. doi:10.1158/0008-5472.Can-13-3116

24. Kong C, Zhan B, Piao C, Zhang Z, Zhu Y, Li Q. Overexpression of UNC5B in bladder cancer cells inhibits proliferation and reduces the volume of transplantation tumors in nude mice. BMC Cancer. 2016/11/15 2016;16(1):892. doi:10.1186/s12885-016-2922-9

25. Lin X, Zhang H, Dai J, et al. TFF3 Contributes to Epithelial-Mesenchymal Transition (EMT) in Papillary Thyroid Carcinoma Cells via the MAPK/ERK Signaling Pathway. J Cancer. 2018;9(23):4430–4439. doi:10.7150/jca.24361

26. Liu J, Kim SY, Shin S, et al. Overexpression of TFF3 is involved in prostate carcinogenesis via blocking mitochondria-mediated apoptosis. Experimental & Molecular Medicine. 2018;50(8):1–11.

27. Yusufu A, Shayimu P, Tuerdi R, Fang C, Wang F, Wang H. TFF3 and TFF1 expression levels are elevated in colorectal cancer and promote the malignant behavior of colon cancer by activating the EMT process. International journal of oncology. 2019;55(4):789–804.

28. Jiale C, Yin S. Mechanism of TFF3-dependent TWIST1 Upregulating TRIB3 to Promote Colorectal Cancer Metastasis. Cancer Research on Prevention and Treatment. 2024;51(2):85–90.

29. Grandin M, Meier M, Delcros JG, et al. Structural Decoding of the Netrin-1/UNC5 Interaction and its Therapeutical Implications in Cancers. Cancer Cell. Feb 8 2016;29(2):173–85. doi:10.1016/j.ccell.2016.01.001

30. Llambi F, Causeret F, Bloch-Gallego E, Mehlen P. Netrin-1 acts as a survival factor via its receptors UNC5H and DCC. The EMBO journal. 2001;20(11):2715–2722. doi:10.1093/emboj/20.11.2715

