## Supplementary Figures for "Netrin-1 regulates colorectal cancer stem cell self-renewal via a TFF3 dependent paracrine survival mechanism"

*Supplementary Figure 1*

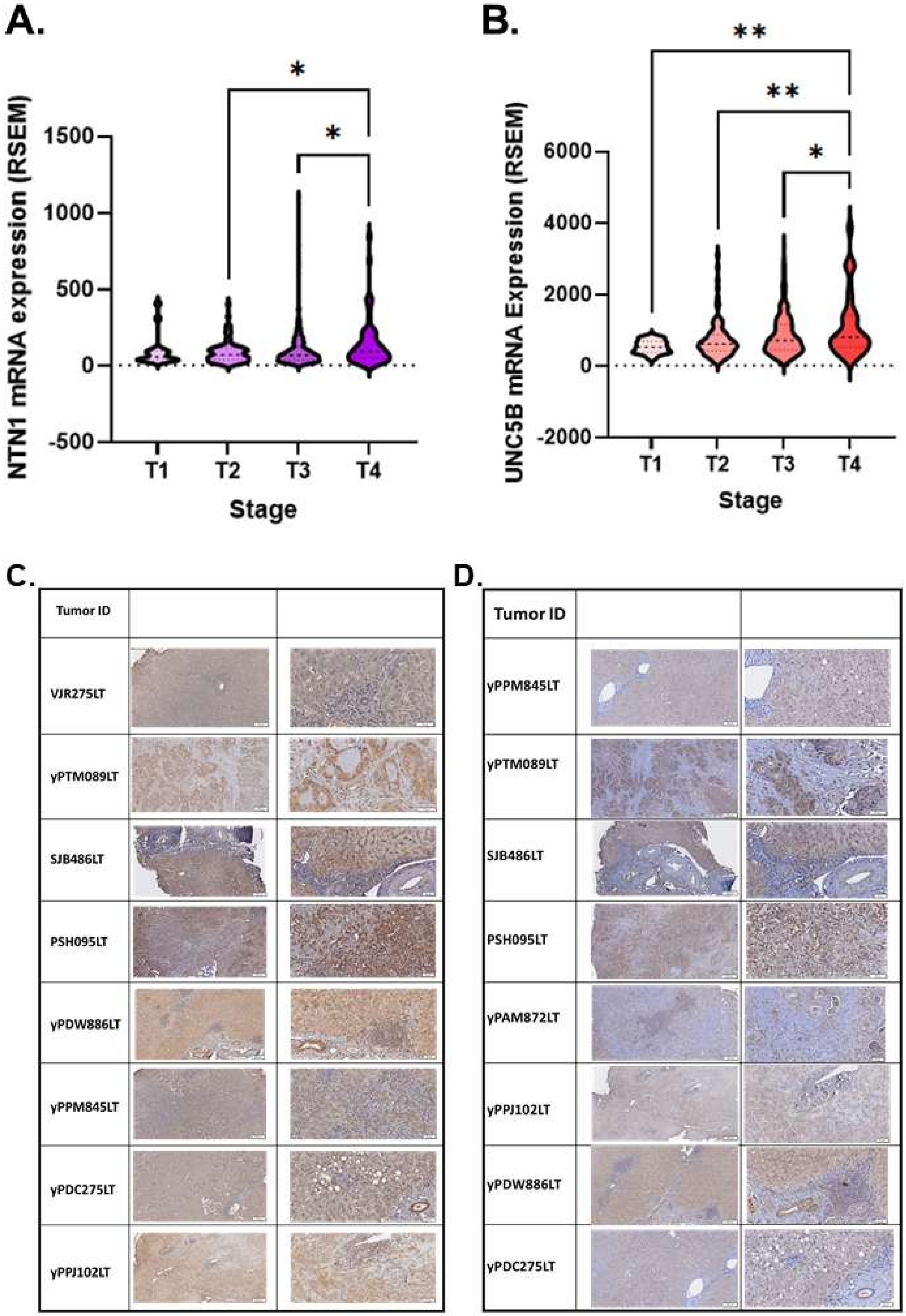

### *Supplementary Figure 2*

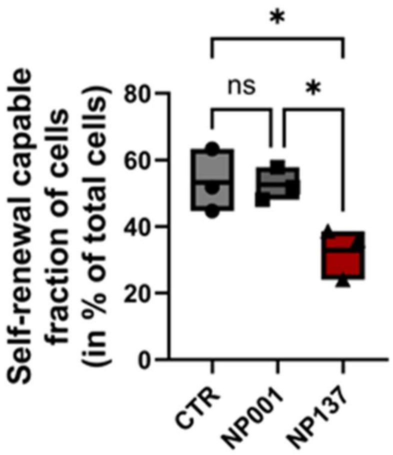


### *Supplementary Figure 3*

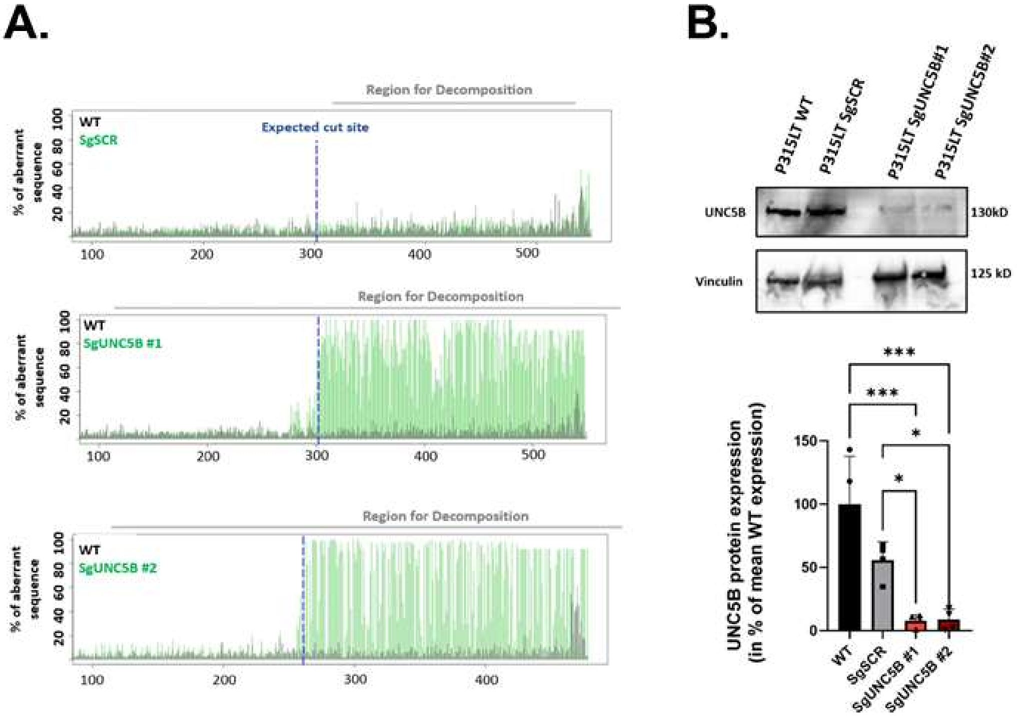

*Supplementary Figure 4*

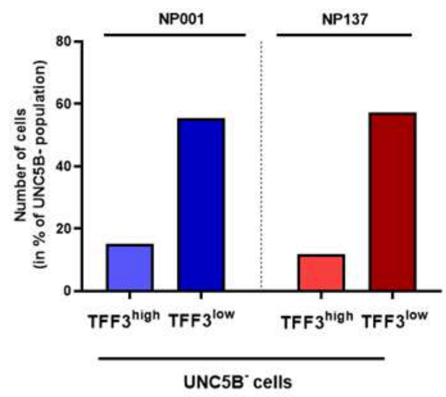


*Supplementary Figure 5*

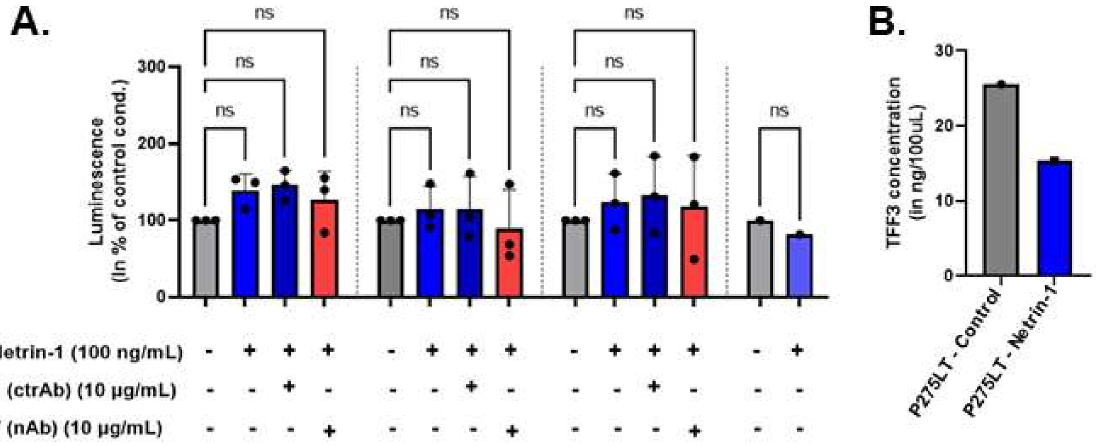


*Supplementary Figure 6*

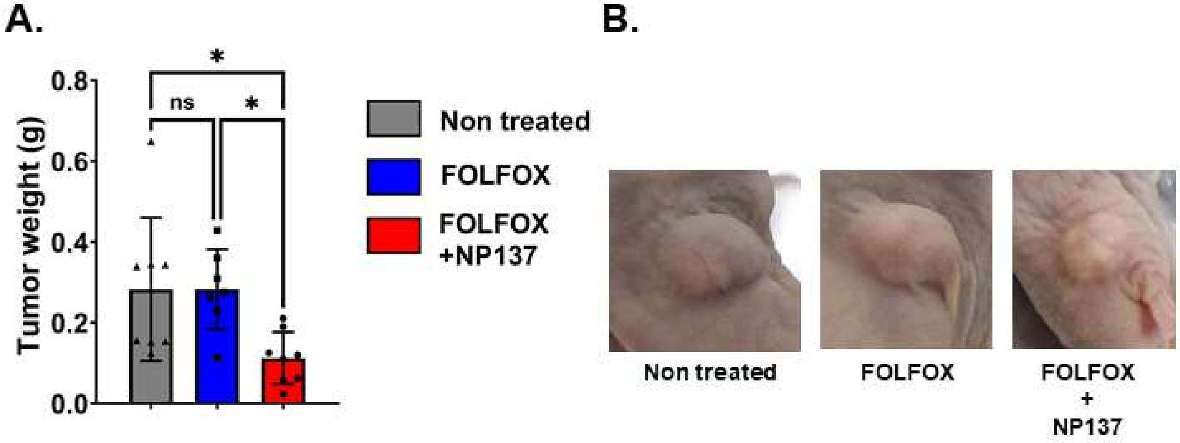
