## Supplementary information for "Netrin-1 regulates colorectal cancer stem cell self-renewal via a TFF3 dependent paracrine survival mechanism"

**Supplementary Figure Legends**

**Supplementary Figure 1 - Analysis of netrin-1 and UNC5B receptor expression in stage CRC tumors - (A-B)** Immunohistochemical Staining of netrin-1 (A) and UNC5B (B) in a panel of Stage IV Colorectal cancer liver metastasis sections (left: scale = 200 um, right: scale = 50 um; Stage IV; **(C)** Expression Profiles of NTN1 mRNA in Colorectal Cancer TCGA according to T stage **(D)** Expression Profiles of UNC5B mRNA in Colorectal Cancer TCGA according to T stages; p Values: * = <0.05 Statistical test: One-Way ANOVA - Tukey's multiple comparisons test; related to Figure 1

**Supplementary Figure 2 - Netrin-1 inhibition is decreasing self-renewal in CRC -** Self-renewal capable fraction of HCT116 cells, either left untreated (Control) or treated with NP001 (10μg/mL) or NP137 (10μg/mL), quantified using ELDA Statistical significance was determined using One-way Anova – Tukey’s multiple comparison test, ns = non-significant, * = p <0.05, ** = p <0.01. N=3; related to Figure 3

**Supplementary Figure 3 - CRISPR Cas9 Invalidation of UNC5B in P315LT mCRC organoid - (A)** Representative TIDE analyses comparing Sanger sequencing traces from wildtype PDOs versus scramble control (sgSCRAMBLE) or CRISPR-edited (SgUNC5B#1 and SgUNC5B#2) organoid **(B)** Western Blot validation of UNC5B protein knockout by looking at the protein level of UNC5B in wildtype, scramble control (sgSCRAMBLE) and CRISPR-edited (SgUNC5B#1 and SgUNC5B#2) in **P315LT**; For Indel spectrum: Green coloured lines represent the percentage of aberrant nucleotides between the sequence trace of the control and the experimental sample, black coloured lines represent noise; Statistical significance was determined using One-way Anova – Tukey’s multiple comparison test, Error bars indicate SD; ns = non-significant, * = p <0.05, *** = p <0.001, N=3; related to Figure 3

**Supplementary Figure 4 – NP137 has no impact on TFF3 expression in UNC5B-negative cells -** Fraction of high-expressing TFF3 and low-expressing TFF3 in UNC5B-positive cells treated with either NP001 (10μg/mL) or NP137 (10μg/mL); High TFF3 expression being defined as > log5 and low expression as < log4; related to Figure 6

**Supplementary Figure 5 - Netrin-1 enhances self-renewal though TFF3** – **(A)** Viability of PDOs (yP190LT; P315LT; P931LT) in the presence or absence of recombinant Netrin-1 (100 ng/mL), NP001 (10μg/mL) and/or NP137 (10μg/mL) ; **(B)** TFF3 secretion in the culture medium of UNC5B and Netrin-1-negative PDOs (VJR275LT) in the presence or absence of recombinant Netrin-1 (100 ng/mL) (n=1), quantified using Cell Titer Glo - 3D (n=3)(n=1 for VJR); ; related to Figure 7

**Supplementary Figure 6 – NP137 treatment enhance FOLFOX treatment *in vivo* (A)** - Tumor weight of engrafted tumors in mice treated with either PBS, FOLFOX or FOLFOX and NP137 at week 10 ; N = 8/group, Error bars indicate SD, Statistical test used : One-way Anova - Tukey's multiple comparisons test;  **(B)** Pictures of representative tumors in each treatment group (Non treated : mean tumor size = 523.94 mm^3^ , tumor size in picture = 499.85 mm^3^ ; FOLFOX treated : mean tumor size = 417.23 mm^3^, tumor size in picture = 435.6675 mm^3^; FOLFOX + NP137 treated: mean tumor size = 183.94 mm^3^, tumor size in picture =200.37 mm^3^);
